# To Improve Protein Sequence Profile Prediction through Image Captioning on Pairwise Residue Distance Map

**DOI:** 10.1101/628917

**Authors:** Sheng Chen, Zhe Sun, Zifeng Liu, Xun Liu, Yutian Chong, Yutong Lu, Huiying Zhao, Yuedong Yang

## Abstract

Protein sequence profile prediction aims to generate multiple sequences from structural information to advance the protein design. Protein sequence profile can be computationally predicted by energy-based method or fragment-based methods. By integrating these methods with neural networks, our previous method, SPIN2 has achieved a sequence recovery rate of 34%. However, SPIN2 employed only one dimensional (1D) structural properties that are not sufficient to represent 3D structures. In this study, we represented 3D structures by 2D maps of pairwise residue distances. and developed a new method (SPROF) to predict protein sequence profile based on an image captioning learning frame. To our best knowledge, this is the first method to employ 2D distance map for predicting protein properties. SPROF achieved 39.8% in sequence recovery of residues on the independent test set, representing a 5.2% improvement over SPIN2. We also found the sequence recovery increased with the number of their neighbored residues in 3D structural space, indicating that our method can effectively learn long range information from the 2D distance map. Thus, such network architecture using 2D distance map is expected to be useful for other 3D structure-based applications, such as binding site prediction, protein function prediction, and protein interaction prediction.

## 1. Introduction

Computational protein design attempts to design a protein sequence that will fold into a predefined structure to perform a desired function. The motivation of studies in this area is not only to supplement, modify, or improve the function of wild-type proteins but also to improve our fundamental comprehension of the relationship between protein sequences, structures, and functions. The past three decades have witnessed significant progress in de novo protein design^1^. More recently, by using Rosetta package, Silva et al. designed potent and selective mimics of anti-cancer drugs IL-2 and IL-15^2^. Such advances have shown the potential to design novel proteins for diagnostic, therapeutic, and industrial purposes. While significant progress has been made, existing protein design approaches remain low success rates^3^. This has led to efforts on building a library of designed sequences, or sequence profiles (sequences randomly generated by specific probabilities of 20 standard amino acids at each site) for guiding experimental screening or directed evolution^4–7^

Typically, protein sequences or sequence profiles can be generated by applying mutations on a random sequence iteratively to minimize its folding free energy with proper optimization algorithm^8–12^. However, the search of global minima is not guaranteed since it’s an NP-hard combinatorial optimization problem^13^. To explore the possibility of more computationally efficient protein design methods, Dai et al. proposed a fragment-based method by searching structurally similar fragments from known protein structures^14, 15^. For a given target protein structure, the sequence profile obtained from structurally similar fragments shows high similarity to its sequence. This fragment-based method is of high computational efficiency but a lack of information on non-local residue interactions (close in three-dimensional structure but not in sequence). Li et al employed a knowledge-based scoring function to compute residue specific energy values according to 3D structures, and integrated them with the profiles derived from fragments into neural networks^16^. The developed SPIN method by training neural network with the local (e.g. fragment-derived) and nonlocal (e.g. energy-based) features achieved a sequence recovery of 30%. Later, the sequence recovery was improved to 33% by using a deep learning method^17^. At the same time, SPIN2^18^, an updated version of SPIN, was also developed by utilizing deep learning network with additional features, slightly improving the sequence recovery to 34.4%. However, all these prediction methods utilized only 1D structural properties that are not sufficient to represent 3D structures.

In order to make a full use of protein 3D structural information, a few studies attempted to input the whole 3D structural information into a 3D-Covolutional Neural Network (3D-CNN) for different biological problems, such as protein-ligand scoring prediction^19^, protein-binding site prediction^20^, side chain conformation prediction^21^, and quality assessment of protein folds^22^. However, it remains challenging to train an accurate 3D-CNN network from the large number of redundant variables involved in the highly sparse 3D matrix with the limited number of 3D structures deposited in the protein data bank (PDB).

On the other hand, it was well known that 3D structure can be alternatively represented by the 2D contact map, which simply shows whether distance of each residue pair is below a threshold (usually 8Å). For example, Skolnick et al. stated that their algorithm was able to successfully fold a small protein even with a small portion of inter-residue contacts^23^. Many recent reports showed that predicted contact map could even produce high-quality 3D protein structures^24^. Moreover, the 2D contact map is an image that can be efficiently processed by modern deep learning techniques, and the prediction from 2D contact maps to sequence profiles is similar to the image captioning problem^25^

There exists to be a few differences with traditional image captioning tasks. First, classical image captioning tasks take input of only a single 2D image, while our inputs include both 2D distance maps and 1D structural features. Second, in image caption scenarios, images are often preprocessed to a fixed size, but our distance maps can’t be resized because each pixel represents exactly one residue pair, and residues far in the sequence might be neighbored in 3D space. Third, the target output of image captioning task is a sentence whose length is irrelevant with input, while our input distance map is of size L × L where L is equal to length of our target output (L × 20).

Inspired by the image captioning tasks, we have designed a novel network architecture coupling bidirectional long short-term memory (BiLSTM) with self-attentional 2D-convolution neural networks (CNN) to predict protein sequence profile, namely SPROF method. The deep neural network can process both 1D structural properties and a 2D distance map reflecting the continuous distances between residue pairs. To our best knowledge, this is the first study to utilize a 2D distance map for structure-based prediction of protein properties. The SPROF method achieved sequence recovery rates of 39.8% on the independent test set, which is significantly higher than 34.6% achieved by the SPIN2 method trained from only 1D structural features. Further analysis indicated that the improvement was mostly contributed by residues most contacted with other residues, suggesting that the inclusion of 2D distance map can efficiently capture long-range contacted information. Therefore, such network architecture to utilize 2D distance map is expected to be useful for other 3D structure-based applications such as binding site prediction, protein function prediction, and protein interaction.

## 2. Materials and Methods

### 2.1 Datasets

Since training deep learning network requires a large number of training samples, we employed the dataset curated in 2017, as used in our previous study^26^. The dataset is consisted of 12450 non-redundant chains with resolution < 2.5Å, R-factor < 1.0, sequence length ≥ 30, and sequence identity ≤ 25% from the cullpdb website. Among them, 11200 chains deposited before Jun 2015 were selected as training set and the remained 1250 were used as an independent test set.

From this dataset, we removed long chains with ≥500 amino acids because the required memory for learning is over the 12GB memory limitation by our used Graphics Processing Unit (GPU) Nvidia GTX 1080 Ti. Finally, we kept a dataset of 7134 chains for training and 922 chains for the test, namely TR7134 and TS922, respectively.

### 2.2 Features Extraction

Our input features include both 1D structural features and 2D distance maps. The 1D structural features include 150 features that are similar to those used in SPIN2^18^. For completeness, we make a brief introduction on the 1D features.

#### 1D structural features

The 1D structural features can be divided into four feature groups: the secondary structures (8), cosine and sine values of backbone angles *ϕ,ψ*,θ, ω, *and τ* (10), local fragment-derived profiles (20), and the global energy features (112), namely GF_SS, GF_AG, GF_FRAG, and GF_ENERGY, respectively. GF_SS are one-hot DSSP codes for eight-state protein secondary structures (C, G, H, I, T, E, B, S). GF_AG are sine and cosine values of 5 backbone angles *ϕ, ψ*, ω, θ, and *τ* at each given position, where *ϕ, ψ*, and ω are three main-chain dihedral angles rotated along *N* – *C_α_, C_α_* – *C*, and *C_i_* – *N*_*i*+1_ bonds, respectively, *τ* is dihedral angle based on four neighboring *C_α_* atoms *C*_*α*_*i*−1__ – *C*_*α_i_*_ – *C*_*α*_*i*+1__ – *C*_*α*_*i*+2__, and θ is angle intervening *C*_*α*_*i*−1__ – *C_α_i__* – *C*_*α*_*i*+1__. GF_FRAG are the probabilities of 20 standard residue types at each position estimated from structurally similar fragments^15^. GF_ENERGY are the interaction energies of 20 standard residue types at a selected position with the rest of the backbone positions occupied by the alanine residue. The energies are computed by using the DFIRE statistical scoring function^27^ based on preferred backbone states-dependent side-chain conformations as defined in the bbdep rotamer library^28^. If one residue type has >6 rotamers, only 6 most frequent rotamers were chosen. We also used the lowest energies among all rotameric states for each residue type. Finally, this generated a total of 112 (=(6+1)×13+(3+1)×4+(2+1)×1+1×2) energy features. Different from SPIN2, we didn’t utilize distances between atoms within the same residue or belonging to neighbored residues since they might include residual information in the force field during the determination of protein 3D structures according to experimental data.

#### 2D distance maps

In addition to the 1D structural features used by SPIN2, we derived an input feature of 2D distance matrix *S* (namely distance map) with its elements

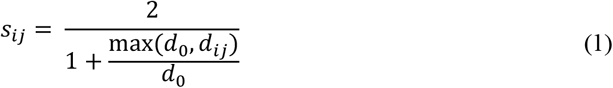

where *d_ij_* is the distance between *C_α_* atoms of residues *i* and *j*, and *d*_0_ was set 4.0 Å, as also used in definition of the SP-score^29^. This conversion of distance ensures a score ranging from 0 to 1, with a good discrimination for distances between 4 and 8 Å. We did not use exactly the same formula as SP-score (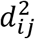 used in SP-score) since it produced slightly worse results (results not shown).

### 2.3 Deep Learning Method

The 2D distance map can be viewed as a special image, with the prediction of protein sequence profile to be producing an image caption for the 2D distance map. Inspired by the image captioning learning architecture, we have designed a deep learning networks coupling RNN and CNN to extract features from 1D and 2D features, respectively. As shown in Figure 1A, a self-attentional ultra-deep residual convolutional Neural Network (ResNet-CNN)^30^ encoded the 2D distance map into a vector representation, which was then concatenated with our 1D structural features, and fed into an RNN module to generate a protein sequence profile.

**Figure 1.**
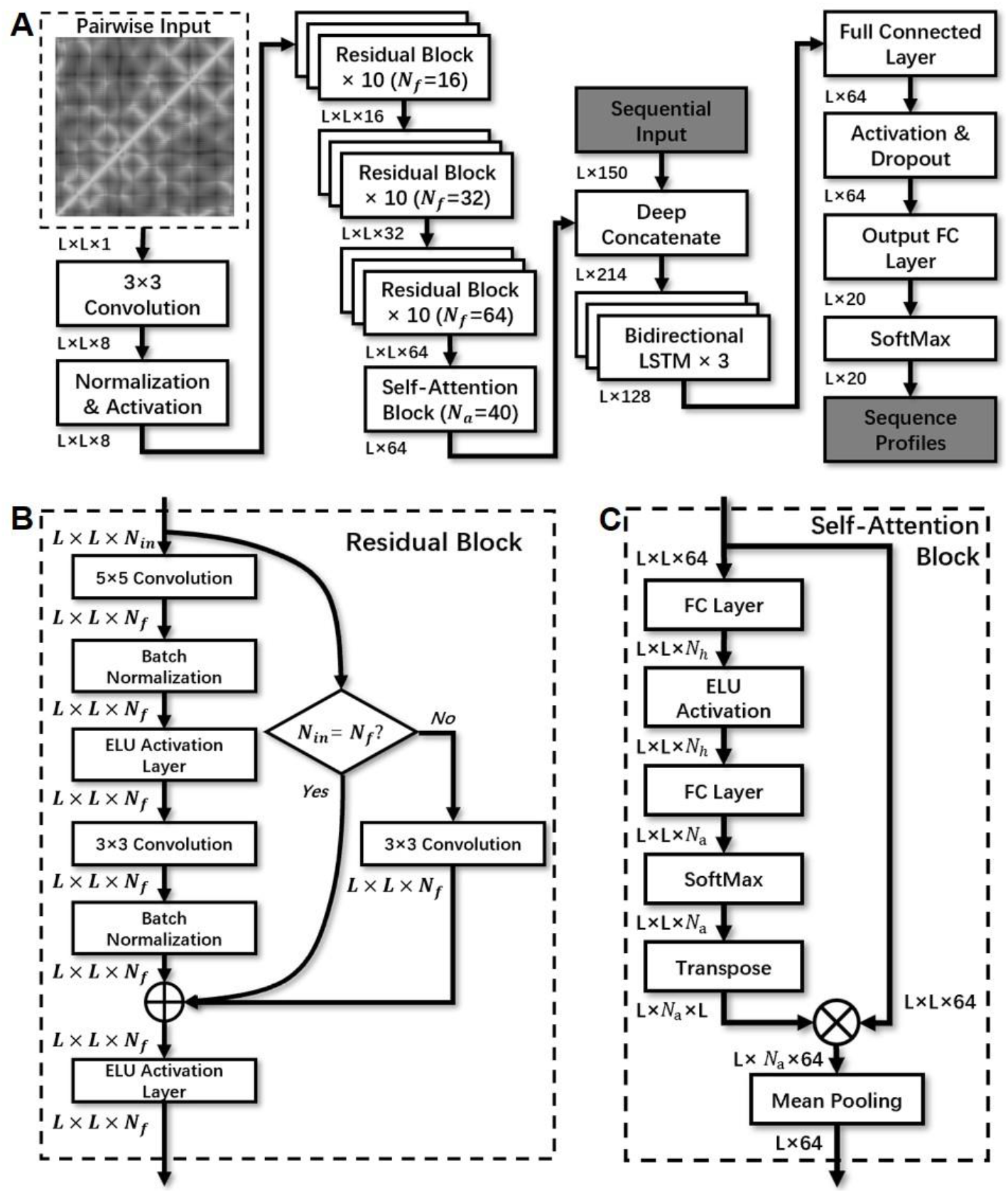
**The neural network layout of SPROF** with (A) illustrating the overall network architecture of SPROF, where L is the length of protein sequence; (B) showing details of the residual block, where *N_in_* is the number of input layers, *N_f_* represents the number of kernels in each convolution layer; (C) indicating the structure of the multi-head self-attention block, where *N_a_* represents the number of parallel attention layers, and *N_h_* is the number of hidden states between two Fully-Connected (FC) layers (chosen as 50 here).

#### CNN module

CNN has demonstrated superior performance in image tasks because of its implementation of shift, scale, and distortion invariance through local receptive fields, shared weights, and sub-sampling. Though the representation depth of CNN is beneficial for the classification^31^, the vanishing gradient problem has become a major obstacle to increasing the depth of CNN. In 2015, He et al. proposed ResNet, an ultra-deep residual Neural Network to solve the vanishing gradient problem by employing shortcut connection between outputs of a current and its previous layers.^30^. Considering that distance map was composed of sparse residue pairs with short distances that play important roles in spatial structures, we also employed a selfattention mechanism to learn a weight tensor for paying more attention to those important regions^32^. Overall, we have chosen the ResNet with self-attention mechanism for encoding the 2D distance map.

#### RNN module

The features from the CNN module and 1D structural features were concatenated together and fed into a bidirectional Long Short-Term Memory Recurrent Neural Network (LSTM-BRNN) to generate the protein sequence profile. Unlike standard feedforward neural networks, RNN retains a state that can represent information from an arbitrarily long context window^33^. However, traditional RNNs have no ability to learn long-range dependencies as a result of gradient vanishing problem. To overcome this problem, Hochreiter and Schmidhuber proposed the LSTM technique^34^ using carefully designed nodes with recurrent edges of fixed unit weight as a solution^35^. Later, RNN with bidirectional LSTM to exploit both preceding and following dependencies was proposed and has been proved to outperform unidirectional ones in framewise phoneme classification^36^. Currently, LSTM-BRNN has been widely used in many bioinformatics studies^37, 38^.

### 2.4 Neural Networks Implementation Details

In SPROF, 2D distance map (L × L with L as the protein length) was encoded by the self-attentional ResNet into sequential tensors (L × 64), and then concatenated with 1D structural features (L × 150). The concatenated features (L × 214) were fed into a bidirectional LSTM to generate a decoded tensor of L× 128. Finally, a series of Fully Connected (FC) layers and activation layers conducted nonlinear-transformations on the output of bidirectional LSTM to obtain a prediction result of (L × 20), which represented possibilities of 20 amino acid types on each sequence position.

Our self-attentional ResNet module is composed of a series of residual blocks (Figure 1B) and a Self-attention Block (Figure 1C).

### Residual block

Our residual block employed Exponential Linear Unit (ELU) as activation layer instead of Rectified Linear Unit (ReLU) used by typical residual blocks. The ELU activation function was shown to be more effective than standard ReLU for learning in the ResNet^39^. Furthermore, before each activation layer, regularization was applied to the network through the use of batch normalization^40^. Considering the limitation by the used GPU memory size (12GB), we employed 30 ResNet blocks (30×2=60 convolution layers) in our final model. It should be noted that a smaller window size with more number of layers is beneficial for performance of CNN^31^. Thus, the kernel size of two convolution layers in each residual block was chosen as 5 × 5 or 3 × 3 respectively.

### Self-attention Block

In order to extract a weight tensor to focus on important regions, we adopted self-attention mechanism with a self-attention block as shown in Figure 1C. This block converts feature-size into L × 64, so that it can be concatenated with sequential features of L × 150. The concatenated feature size is L × 214.

### To Handle Variable Length Inputs

Different from general image tasks that often preprocess images to the same size, protein sequence profile prediction has to handle proteins of variable sizes. Therefore, we had to design a CNN that could process inputs of variable sizes and ensure the output have a size equaling to its input. Finally, our neural networks don’t have pooling layers as CNN networks often do, and the output of the last residual block remains the same value (L) for width and height.

### Bidirectional LSTM

The input of our bidirectional LSTM is in size of L × 214. The bidirectional LSTM module consists of 3 layers of bidirectional LSTM. In each layer, there are two independent LSTM representing two directions, respectively. Our LSTM cells consist of 64 one-cell memory blocks, culminating in 128 hidden states for each bidirectional LSTM layer.

### Linear Layers

Our linear layers are fully connected. The first FC layer consist of 64 nodes plus a bias node with an ELU activation. The FC output layer has 20 output neurons and a sigmoid activation to convert the output into a likelihood of each amino acid type at each position (L × 20).

### Tools

We trained our model in the framework of Facebook’s PyTorch library (v0.4.0), which enables us to accelerate the model training on an Nvidia GeForce GTX 1080 Graphics Processing Unit (GPU). It has been shown that the use of a GPU for training a neural network can speed up by a factor up to 20^41^.

### Optimization algorithm and Dropout

Our model was trained with cross entropy as the loss function and ADAM algorithm for optimization^42^. ADAM optimization algorithm is generally considered to be robust for the selection of hyperparameters and converges more quickly than the traditionally-used Stochastic Gradient Descent (SGD). We used a learning rate of 0.001 in this study. Furthermore, a 50% dropout rate was adopted at the output of the fully-connected layer during training to reduce overfitting^43^.

### Hyperparameters Tuning by the Cross Validation

The architecture and hyperparameters were optimized by the 5-fold cross validation, where the training set was randomly divided into five different subsets. Each time four of these subsets were used to train a model and the left one was used for the test. This process was repeated for five times so that all five subsets were tested exactly once, and the average accuracy over five tests was used for the overall performance. With the hyperparameters achieving the best performance, the final model was trained on the whole training set, and tested on the independent test set.

### Evaluation

We evaluated the performance by the native sequence recovery that is the percentage of residues that were correctly predicted. A residue was considered to be correctly predicted if the wild type residue type has the highest value in the predicted profile for 20 residue type at the position.

In addition, we also evaluated the performance for different types of residues, we calculated precision and recall for residue R as

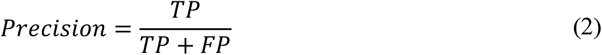

and

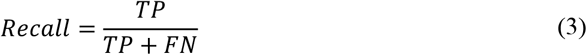

where TP is the number of correctly predicted residues for type *R*, FP is the number of residues wrongly predicted as *R*, and FN is the number of incorrectly predicted residues of wild-type R.

## 3. Results

### 3.1 Model Selection and Feature Importance

Table 1 illustrates the performance of SPROF and its variants with different network architectures. SPROF achieved sequence recovery of 39.9% and 39.8% for the 5-fold crossvalidation (CV) and independent test, respectively. The consistent results indicate the robustness of the SPROF method. An exclusion of the self-attention block (SPROF-noAtt) caused a decrease of 0.9% in the CV and 1.2% in the independent test. The removal of RNN module and 1D structural features (SPROF-RNN) decreases the sequence recovery to 36.1% in the CV. The slight higher recovery percentage (36.3%) in the test set than the CV should result from random fluctuations. The greatest drop was from SPROF-CNN that excluded the CNN module and 2D distance map. This caused 6% and 6.6% drop of native sequence recoveries in the training set and independent test, respectively. The results demonstrate the benefits of utilizing the distance map features and image captioning method.

**Table 1.** Native sequence recovery rates achieved by the SPROF and its variants for 5-fold cross validation and independent test.

| Model | TR7134 | TS922 |
| --- | --- | --- |
| SPROF <sup>a</sup> | 39.9% | 39.8% |
| SPROF-noAtt <sup>b</sup> | 39.0% | 38.6% |
| SPROF-CNN <sup>c</sup> | 36.1% | 36.3% |
| SPROF-RNN <sup>d</sup> | 33.9% | 33.2% |

<sup>a</sup> The best performed model, detailed shown in Figure 1.
<sup>b</sup> SPROF without self-attention block.
<sup>c</sup> Using the CNN module without bidirectional LSTM module or input of 1D structural features.
<sup>d</sup> Using the RNN module without self-attentional ResNet module or input of 2D distance maps.

It is of interests to see which type of features made the greatest contribution in the prediction. We excluded each type of features one-by-one to obtain five different feature sets for model training, and then compared the performance of each model. Table 2 shows the sequence recovery of these five models in the independent test set. As expected, 2D distance map features contributed the most in the sequence recovery (contributing 6.6% on independent test), followed by energy-based features (1.8%) that made the highest contribution in the SPIN2. The exclusion of fragment-based features made overall sequence recovery 0.7% lower, and the exclusion of secondary structure features or backbone torsion angles features also marginally decreased the overall sequence recovery (0.3% and 0.2%, respectively). These results highlight the importance of distance map in our prediction model, which inspired us to employ distance map features on other 3D structure-based applications in future.

**Table 2.** The comparison of sequence recovery after excluding one feature group from SPROF.

| **Feature Excluded** | **TR7134** | **TS922** |
| --- | --- | --- |
| SPROF | 39.9% | 39.8% |
| - distance map | 33.9% | 33.2% |
| - GF_ENERGY | 37.8% | 38.0% |
| - GF_FRAG | 38.7% | 39.1% |
| - GF_SS | 39.6% | 39.5% |
| - GF_AG | 39.7% | 39.6% |

### 3.2 Comparison with other methods

We further made direct comparison with SPIN2 on the test set TS922. As shown in Table 3, there is over 5% consistent improvement from SPIN2 to SPROF in the native sequence recovery for both top 1 and top 2 matches. Since Wang’s method^17^ is not available online, we can’t make a direct comparison. According to published results, SPIN2 and Wang’s method should be close because SPIN2 is over 4% higher than SPIN, while Wang’s method is about 3% higher than SPIN.

**Table 3.** Overall sequence recovery comparison between SPIN2 and SPROF

| Method | TS922 <sup>a</sup> | Top2 Match <sup>b</sup> |
| --- | --- | --- |
| SPROF | 39.8% | 54.8% |
| SPIN2 | 34.6% | 49.4% |

<sup>a</sup> The percentage of mapped residues with the predicted on TS922.
<sup>b</sup> Match of wild-type sequence to one of top 2 predicted residue types on TS922.

We compared the performance of SPROF, SPROF-CNN, SPROF-RNN, and SPIN2 for proteins on TS922 with different lengths. As shown in Figure 2A, SPROF consistently outperformed SPIN2 in all intervals and SPROF-CNN model is somewhere in between. SPROF-RNN model is less accurate than SPIN2, likely because SPROF-RNN model excluded partial features employed by SPIN2. A direct comparison of the sequence recovery rates (Figure 2B) suggests that SPROF is significantly better than SPIN2 (P-value<10^−99^) according to the pairwise t-test, where SPROF outperformed SPIN2 for 815 proteins, worse for 76 proteins, and tied for the remained 31 proteins.

**Figure 2.**
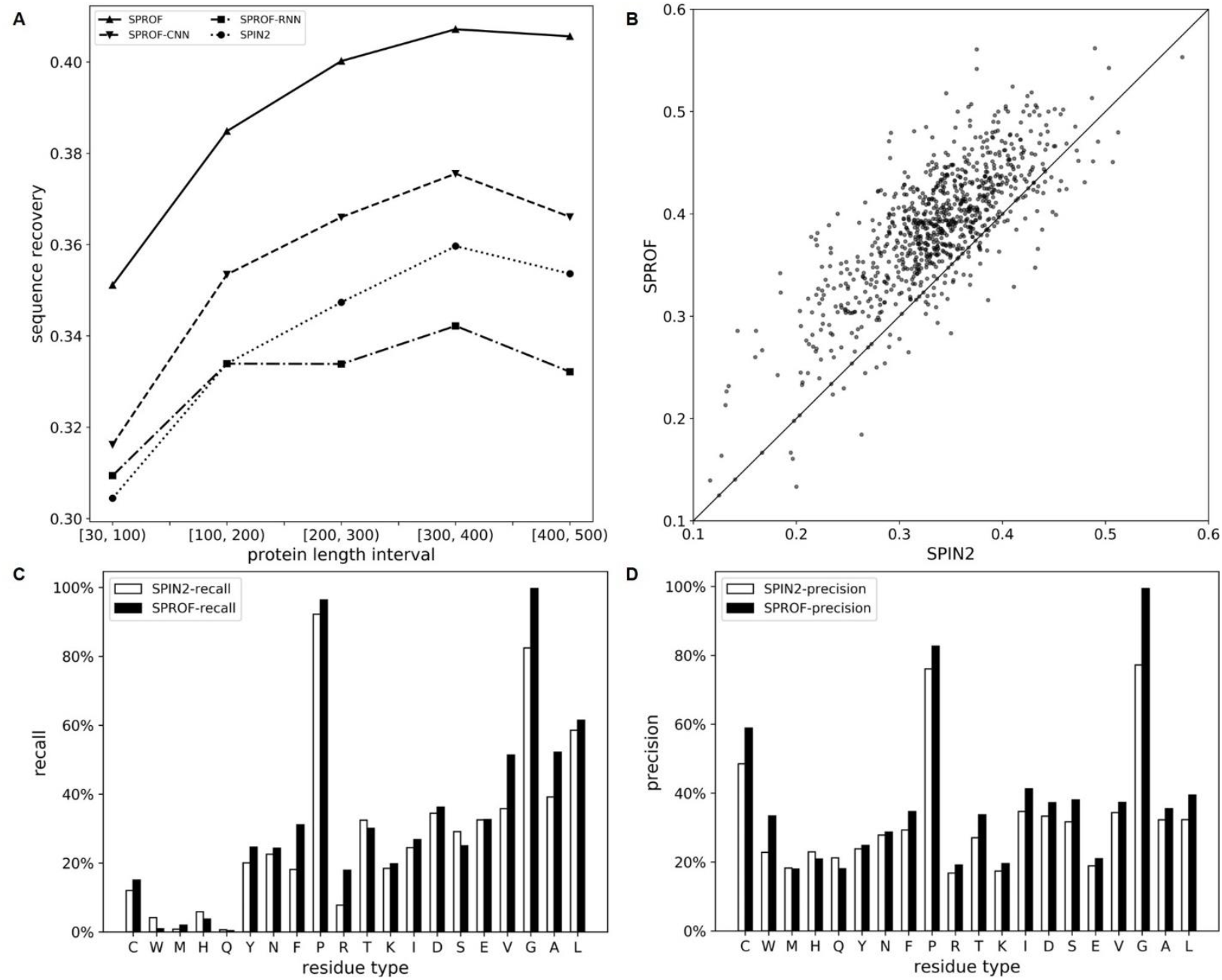
(A) The average sequence recovery rates of different length intervals by four methods; (B) the sequence recovery for each chain in TS922 by SPROF and SPIN2; (C) the recall and (D) precision for different amino acids residue types by SPIN2 and SPROF over TS922.

For a given residue type, we compared the recall and precision score of SPROF and SPIN2 on TS922, as shown in Figure 2C and Figure 2D, respectively. SPROF outperformed SPIN2 in 15 (75% of 20) amino acids types for recall and 17 (85% of 20) for precision.

To explore why SPROF outperformed SPIN2, we plotted the prediction accuracy of residues as a function of their contact number for different methods. The contact number was defined as the number of neighboring C_α_ atoms no farther than 13 Å from a given C_α_ atom. As shown in Figure 3A, SPROF and SPROF-CNN show an increase of prediction accuracy with the addition of neighbored residues. By comparison, SPROF-RNN and SPIN2 without using 2D distance map show a close to flat performances for all residues. This comparison indicated that the inclusion of 2D distance map helps to capture information of residues contacted in 3D structure.

**Figure 3.**
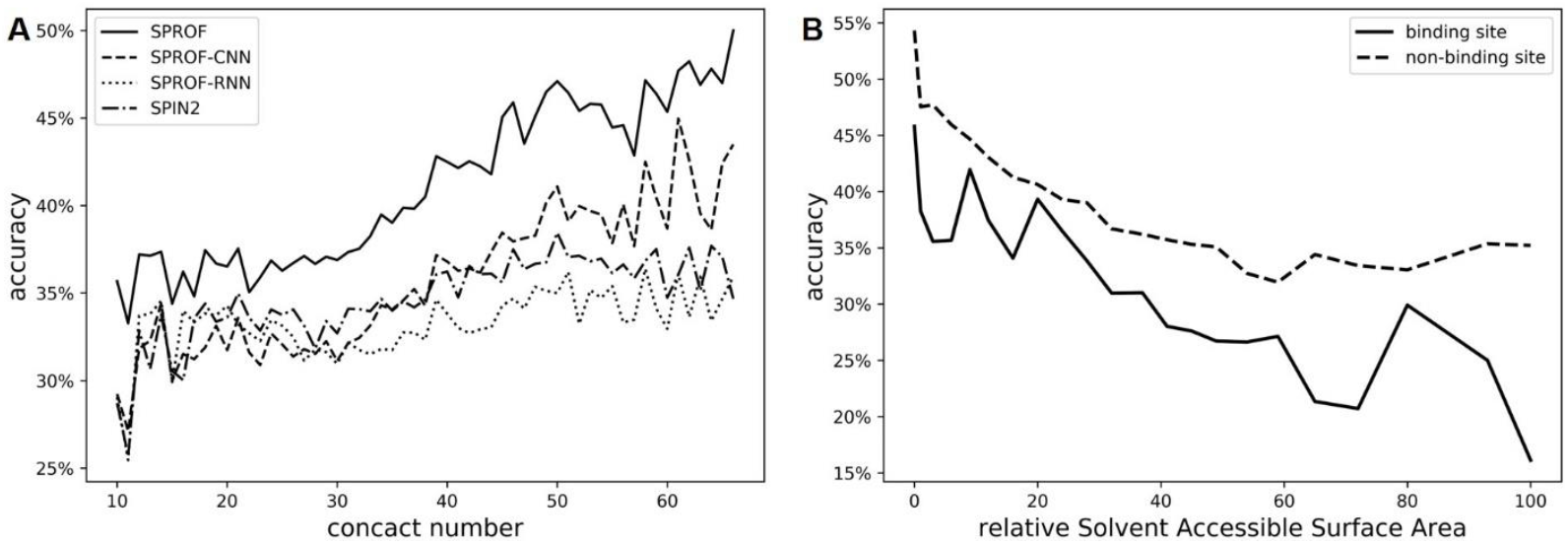
(A) The accuracy for residues as a function of their contact numbers for SPROF and SPIN2; (B) the line plot of prediction accuracy for residues of different rASA intervals and binding or non-binding site on 357 chains (overlap with BioLip) of TS922.

We also compared the prediction accuracies between binding and non-binding sites. By mapping the proteins of TS922 to those defined in BioLip^44^, we generated a dataset of 357 chains dataset. As shown in Figure 3B, the prediction accuracies of residues decrease with the relative solvent surface area, and the accuracy of binding residues is consistently lower than that of non-binding residues. This is as expected because buried residues maintain 3D spatial structures, and binding residues are evolved mainly for protein function, and not necessary to be optimized for 3D structure.

### 3.3 Case study

To illustrate our method, we chose the precorrin-6A reductase cobK (pdbID: 5c4n chain D) for comparisons of methods. The protein chain contains 8 helical and 12 beta sheet fragments, totally 244 amino acids. For a clear look of the predicted sequence profile, we plotted sequence logos for fragment of residue index 75-104, the red part in the Figure 4A. SPROF and SPIN2 achieved accuracies of 60% and 26.7% for the fragment, respectively. As shown in the Figure 4C and D, SPROF has made correct prediction for 11 amino acids (red amino acids in Figure 4E) that are not correctly predicted by SPIN2. A deep look indicates most of the amino acids in the list are hydrophobic (6 Alanine and 1 Valine). This result is consistent with our expectation because our method is better for predicting most contacted residues that are frequently hydrophobic amino acids. SPROF only misses one prediction (No. 96) that is correctly predicted by SPIN2. On this position, the native amino acid (Threonine) ranked the 3^rd^ by our prediction.

**Figure 4.**
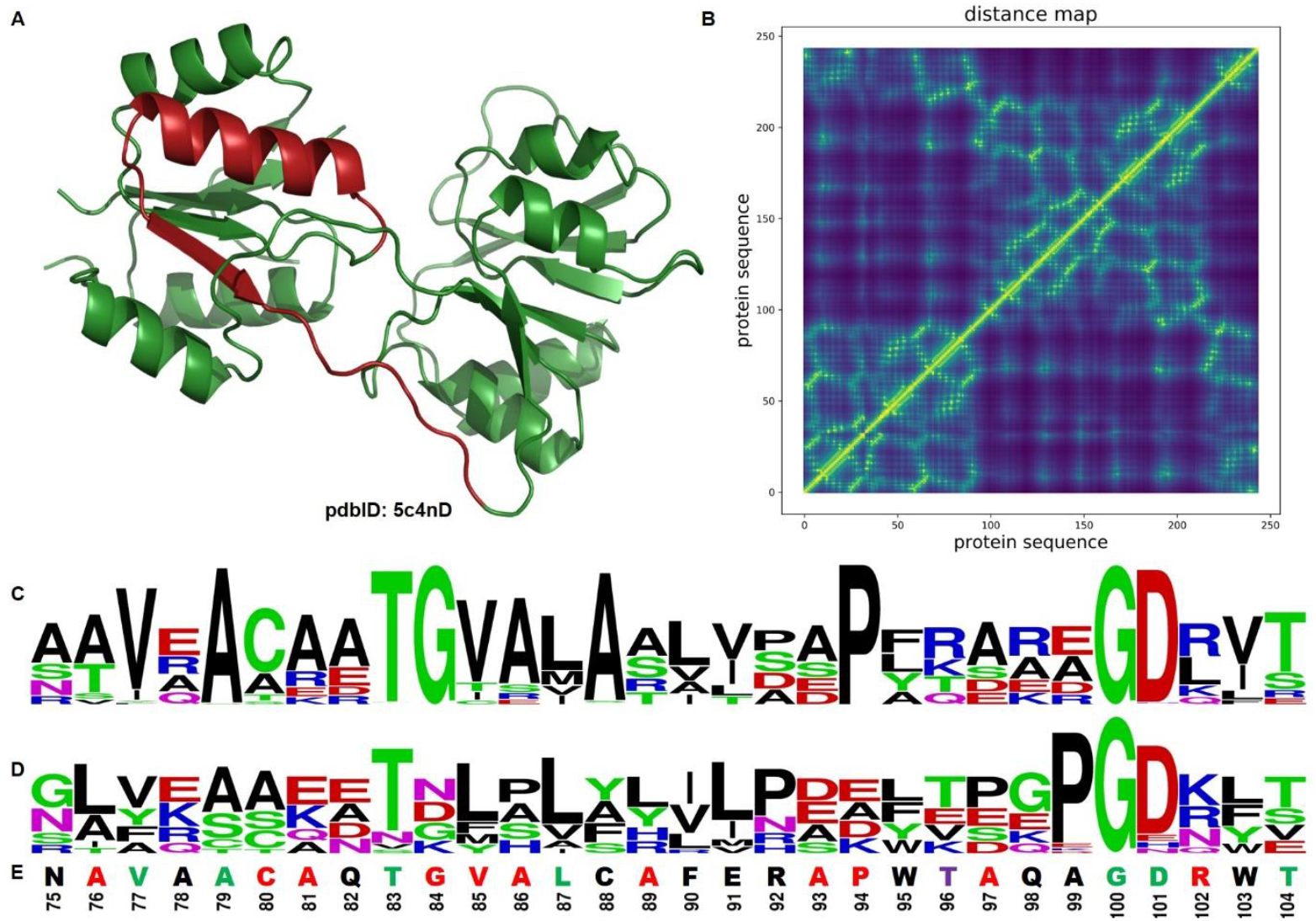
The (A) 3D structure, (B) Distance map, and sequence logo generated by (C) SPROF and (D) SPIN2 for the precorrin-6A reductase cobK (pdbID: 5c4nD). For a clear look, only fragment 75-104 (red in the 3D structure) was shown in the sequence logo generated by SPIN2 and SPROF. The wild-type sequence and indexes were provided in (E) with red, purple, green, and black for correct prediction of amino acids by SPORF only, SPIN2 only, both methods, and none, respectively.

## 4. Conclusions and Discussions

This study highlights the power of applying image captioning method on 2D distance map for protein sequence profile prediction. We proposed a protein sequence profile prediction method SPROF which combined recurrent neural network, convolution neural network, and attention mechanism. SPROF has improved the native sequence recovery from 34.6% (previous method SPIN2) to 39.8% on our independent test set. The improvement is consistent regardless of proteins lengths, test sets (cross-validation and independent test), evaluation metrics (top1 and top2 matches, precision and recall score), or types of amino acids. We also trained a model by using only 1D structural features, which is significantly lower than SPROF with inclusion of 2D distance map. This is reasonable because distance maps are capable of indicating the 3D structural information of proteins. Designed by the inspiration of image captioning method, SPROF is capable of extracting these 3D structural information and thus obtains higher accuracy for sequence prediction. Therefore, such network architecture applying image captioning method on 2D distance map is expected to be useful for other 3D structure-based applications, such as binding site prediction, protein function prediction, and protein interaction prediction.

We have shown there is a significant difference of the native residue recovery between binding and non-binding residues, such profile may be employed for discrimination of functional residues. In addition, the generated sequence profiles have been proven beneficial for improving existing protein design and fold recognition techniques studies^14, 45^, so our improved prediction of the sequence profile could advance the applications in future.

## Author Contributions

S.C., Z.S., Z.L., X.L., Y.C., Y.L., H.Z. and Y.Y. designed the method; S.C. and Z.S. developed and implemented methods and produced results; S.C., Z.S, H.Z. and Y. Y. wrote the manuscript; all authors reviewed the manuscript.

## Notes

The authors declare no competing financial interest

## Acknowledgements

This work has been supported by the in part by the National Natural Science Foundation of China (61772566, U1611261 and 81801132) and the program for Guangdong Introducing Innovative and Entrepreneurial Teams (2016ZT06D211).

---

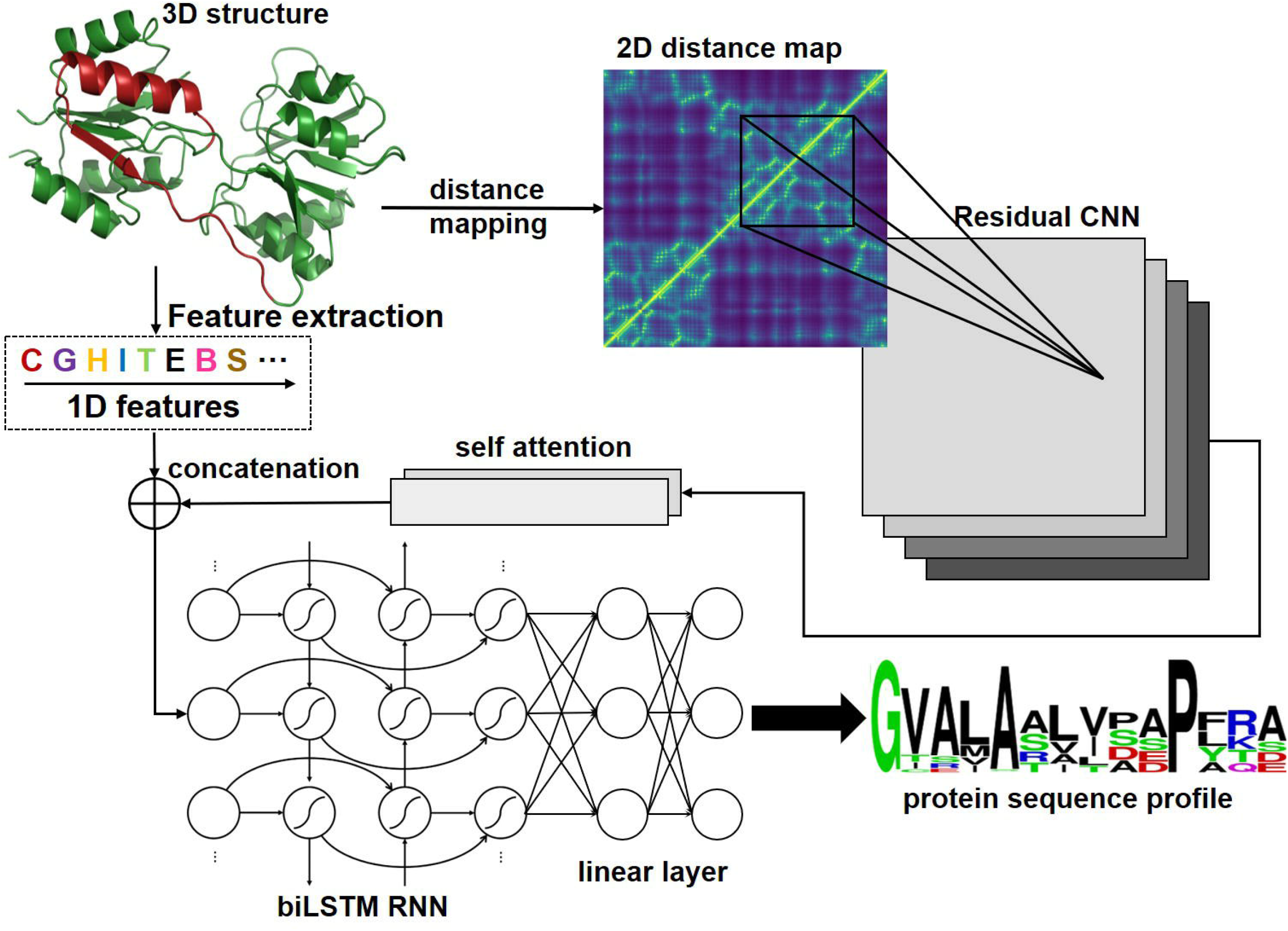

## References

1. Liu, H.; Chen, Q., Computational protein design for given backbone: recent progresses in general method-related aspects. Current opinion in structural biology 2016, 39, 89–95.

2. Silva, D. A.; Yu, S.; Ulge, U. Y.; Spangler, J. B.; Jude, K. M.; Labao-Almeida, C.; Ali, L. R.; Quijano-Rubio, A.; Ruterbusch, M.; Leung, I.; Biary, T.; Crowley, S. J.; Marcos, E.; Walkey, C. D.; Weitzner, B. D.; Pardo-Avila, F.; Castellanos, J.; Carter, L.; Stewart, L.; Riddell, S. R.; Pepper, M.; Bernardes, G. J. L.; Dougan, M.; Garcia, K. C.; Baker, D., De novo design of potent and selective mimics of IL-2 and IL-15. Nature 2019, 565, 186–191.

3. Li, Z.; Yang, Y.; Zhan, J.; Dai, L.; Zhou, Y., Energy Functions in De Novo Protein Design: Current Challenges and Future Prospects. Annual Review of Biophysics 2013, 42, 315–335.

4. Treynor, T. P.; Vizcarra, C. L.; Nedelcu, D.; Mayo, S. L., Computationally designed libraries of fluorescent proteins evaluated by preservation and diversity of function. Proceedings of the National Academy of Sciences 2007, 104, 48–53.

5. Guntas, G.; Purbeck, C.; Kuhlman, B., Engineering a protein–protein interface using a computationally designed library. Proceedings of the National Academy of Sciences 2010, 107, 19296–19301.

6. Allen, B. D.; Nisthal, A.; Mayo, S. L., Experimental library screening demonstrates the successful application of computational protein design to large structural ensembles. Proceedings of the National Academy of Sciences 2010, 107, 19838–19843.

7. Hayes, R. J.; Bentzien, J.; Ary, M. L.; Hwang, M. Y.; Jacinto, J. M.; Vielmetter, J.; Kundu, A.; Dahiyat, B. I., Combining computational and experimental screening for rapid optimization of protein properties. Proceedings of the National Academy of Sciences 2002, 99, 15926–15931.

8. Dahiyat, B. I.; Mayo, S. L., De Novo Protein Design: Fully Automated Sequence Selection. Science 1997, 278, 82–87.

9. Dantas, G.; Kuhlman, B.; Callender, D.; Wong, M.; Baker, D., A Large Scale Test of Computational Protein Design: Folding and Stability of Nine Completely Redesigned Globular Proteins. Journal of Molecular Biology 2003, 332, 449–460.

10. Lippow, S. M.; Tidor, B., Progress in computational protein design. Current opinion in biotechnology 2007, 18, 305–311.

11. Liu, Y.; Kuhlman, B., RosettaDesign server for protein design. Nucleic Acids Res 2006, 34, W235–8.

12. Regan, L.; DeGrado, W. F., Characterization of a helical protein designed from first principles. Science 1988, 241, 976–978.

13. Pierce, N. A.; Winfree, E., Protein design is NP-hard. Protein engineering 2002, 15, 779–782.

14. Zhou, H.; Zhou, Y., Fold recognition by combining sequence profiles derived from evolution and from depth - dependent structural alignment of fragments. Proteins: Structure, Function, and Bioinformatics 2005, 58, 321–328.

15. Dai, L.; Yang, Y.; Kim, H. R.; Zhou, Y., Improving computational protein design by using structure - derived sequence profile. Proteins: Structure, Function, and Bioinformatics 2010, 78, 2338–2348.

16. Li, Z.; Yang, Y.; Faraggi, E.; Zhan, J.; Zhou, Y., Direct prediction of profiles of sequences compatible with a protein structure by neural networks with fragment-based local and energy-based nonlocal profiles. Proteins 2014, 82, 2565–73.

17. Wang, J.; Cao, H.; Zhang, J. Z.; Qi, Y., Computational protein design with deep learning neural networks. Scientific reports 2018, 8, 6349.

18. O’Connell, J.; Li, Z.; Hanson, J.; Heffernan, R.; Lyons, J.; Paliwal, K.; Dehzangi, A.; Yang, Y.; Zhou, Y., SPIN2: Predicting sequence profiles from protein structures using deep neural networks. Proteins: Structure, Function, and Bioinformatics 2018, 86, 629–633.

19. Ragoza, M.; Hochuli, J.; Idrobo, E.; Sunseri, J.; Koes, D. R., Protein–ligand scoring with convolutional neural networks. Journal of chemical information and modeling 2017, 57, 942–957.

20. Jiménez, J.; Doerr, S.; Martínez-Rosell, G.; Rose, A.; De Fabritiis, G., DeepSite: protein-binding site predictor using 3D-convolutional neural networks. Bioinformatics 2017, 33, 3036–3042.

21. Liu, K.; Sun, X.; Ma, J.; Zhou, Z.; Dong, Q.; Peng, S.; Wu, J.; Tan, S.; Blobel, G.; Fan, J., Prediction of amino acid side chain conformation using a deep neural network. arXiv preprint arXiv: 1707.08381 2017.

22. Derevyanko, G.; Grudinin, S.; Bengio, Y.; Lamoureux, G., Deep convolutional networks for quality assessment of protein folds. Bioinformatics 2018, 34, 4046–4053.

23. Skolnick, J.; Kolinski, A.; Ortiz, A. R., MONSSTER: a method for folding globular proteins with a small number of distance restraints. Journal of molecular biology 1997, 265, 217–241.

24. Adhikari, B.; Cheng, J., CONFOLD2: improved contact-driven ab initio protein structure modeling. BMC Bioinformatics 2018, 19, 22.

25. Vinyals, O.; Toshev, A.; Bengio, S.; Erhan, D. Show and tell: A neural image caption generator. In Proceedings of the IEEE conference on computer vision and pattern recognition, 2015; 2015; pp 3156–3164.

26. Hanson, J.; Paliwal, K.; Litfin, T.; Yang, Y.; Zhou, Y., Accurate prediction of protein contact maps by coupling residual two-dimensional bidirectional long short-term memory with convolutional neural networks. Bioinformatics 2018, 34, 4039–4045.

27. Yang, Y.; Zhou, Y., Ab initio folding of terminal segments with secondary structures reveals the fine difference between two closely related all-atom statistical energy functions. Protein Science: A Publication of the Protein Society 2008, 17, 1212.

28. Dunbrack Jr, R. L.; Cohen, F. E., Bayesian statistical analysis of protein side - chain rotamer preferences. Protein Science 1997, 6, 1661–1681.

29. Yang, Y.; Zhan, J.; Zhao, H.; Zhou, Y., A new size - independent score for pairwise protein structure alignment and its application to structure classification and nucleic - acid binding prediction. Proteins: Structure, Function, and Bioinformatics 2012, 80, 2080–2088.

30. He, K.; Zhang, X.; Ren, S.; Sun, J. Deep residual learning for image recognition. In Proceedings of the IEEE conference on computer vision and pattern recognition, 2016; 2016; pp 770–778.

31. Simonyan, K.; Zisserman, A., Very deep convolutional networks for large-scale image recognition. arXiv preprint arXiv:1409.1556 2014.

32. Lin, Z.; Feng, M.; Santos, C. N. d.; Yu, M.; Xiang, B.; Zhou, B.; Bengio, Y., A structured self-attentive sentence embedding. arXiv preprint arXiv:1703.03130 2017.

33. Mikolov, T.; Karafiát, M.; Burget, L.; Černocký, J.; Khudanpur, S. Recurrent neural network based language model. In Eleventh Annual Conference of the International Speech Communication Association, 2010; 2010.

34. Hochreiter, S.; Schmidhuber, J., Long short-term memory. Neural computation 1997, 9, 1735–1780.

35. Lipton, Z. C.; Berkowitz, J.; Elkan, C., A critical review of recurrent neural networks for sequence learning. arXiv preprint arXiv:1506.00019 2015.

36. Graves, A.; Schmidhuber, J., Framewise phoneme classification with bidirectional LSTM and other neural network architectures. Neural Networks 2005, 18, 602–610.

37. Heffernan, R.; Yang, Y.; Paliwal, K.; Zhou, Y., Capturing non-local interactions by long short-term memory bidirectional recurrent neural networks for improving prediction of protein secondary structure, backbone angles, contact numbers and solvent accessibility. Bioinformatics 2017, 33, 2842–2849.

38. Hanson, J.; Yang, Y.; Paliwal, K.; Zhou, Y., Improving protein disorder prediction by deep bidirectional long short-term memory recurrent neural networks. Bioinformatics 2017, 33, 685–692.

39. Shah, A.; Kadam, E.; Shah, H.; Shinde, S.; Shingade, S. Deep residual networks with exponential linear unit. In Proceedings of the Third International Symposium on Computer Vision and the Internet, 2016; ACM: 2016; pp 59–65.

40. Ioffe, S.; Szegedy, C., Batch normalization: Accelerating deep network training by reducing internal covariate shift. arXiv preprint arXiv:1502.03167 2015.

41. Oh, K.-S.; Jung, K., GPU implementation of neural networks. Pattern Recognition 2004, 37, 1311–1314.

42. Kingma, D. P.; Ba, J., Adam: A method for stochastic optimization. arXiv preprint arXiv:1412.6980 2014.

43. Srivastava, N.; Hinton, G.; Krizhevsky, A.; Sutskever, I.; Salakhutdinov, R., Dropout: a simple way to prevent neural networks from overfitting. The Journal of Machine Learning Research 2014, 15, 1929–1958.

44. Yang, J.; Roy, A.; Zhang, Y., BioLiP: a semi-manually curated database for biologically relevant ligand–protein interactions. Nucleic acids research 2012, 41, D1096–D1103.

45. Schmidt am Busch, M.; Mignon, D.; Simonson, T., Computational protein design as a tool for fold recognition. Proteins: Structure, Function, and Bioinformatics 2009, 77, 139–158.

